# Ancient human mitochondrial genomes encode antimicrobial peptides

**DOI:** 10.64898/2026.08.19.745802

**Authors:** Marcelo D. T. Torres, Abdullah Ali, Hyun-Su Lee, Fangping Wan, Cesar de la Fuente-Nunez

## Abstract

Mitochondria are bacteria-derived organelles that coordinate major innate immune pathways, but whether mitochondrial genomes themselves encode direct antimicrobial functions is underexplored. Here, we test the hypothesis that ancient human mitochondrial DNA is not only an evolutionary record, but also contains encrypted peptide sequences with the capacity to contribute to host defense. Because mitochondria descend from a bacterial endosymbiont and retain bacterial-like molecular features, we reasoned that their compact genomes might preserve sequence fragments capable of engaging bacterial-like membranes or bacterial physiology. We mined 2,025 ancient human mitochondrial genomes using a computational pipeline that couples ORF extraction and deep learning, identifying 65 candidate peptides that we term mitochondrins. We synthesized 38 candidates and experimentally validated 14 as antimicrobials against clinically relevant Gram-negative and Gram-positive bacteria. Active mitochondrins were not defined by length, charge, or helicity alone; instead, potency depended on precise hydrophobic-cationic patterning, with nested peptide families revealing how single motif-level changes can switch activity on or off. Mechanistic assays showed that mitochondrins span multiple antibacterial modalities, from strong membrane disruption to potent activity with limited membrane perturbation, suggesting noncanonical or multi-step killing mechanisms. Several peptides displayed low cytotoxicity toward human cells, and one representative mitochondrin reduced bacterial burden in a murine skin abscess model. These findings provide a biochemical basis for the hypothesis that mitochondrial genomes may contribute to innate immunity.

## Introduction

Mitochondria occupy a distinctive position in human immunity. Best known as bioenergetic organelles, they descend from a once free-living bacterium and retain bacterial hallmarks in genome organization, gene expression, membrane biology, and oxidative phosphorylation^1–3^. They are also central regulators of innate immune signaling^4^. Yet this convergence of bacterial ancestry and immune function raises a largely unexplored question: can the mitochondrial genome itself encode molecules with direct antimicrobial activity, thereby contributing to host defense?

This possibility is biologically plausible for two reasons. First, mitochondrial coding space can give rise to bioactive molecules beyond the canonical oxidative-phosphorylation machinery: mitochondrial-derived peptides such as humanin and MOTS-c have cytoprotective and immunomodulatory functions^5–7^. Second, endosymbiosis created a long-standing interface between a bacteria-derived compartment and its host, in which control of membrane integrity, symbiont physiology, and interactions with competing microbes would have been central. We therefore hypothesized that this evolutionary history may have preserved short peptide motifs with latent antibacterial activity.

Short cationic and amphipathic peptides are plausible effectors for such a role because similar molecules act broadly as antimicrobials and host-defense factors. We do not assume that mitochondrial sequences evolved as antibiotics in the modern clinical sense. Rather, fragments embedded within mitochondrial coding space may have originated, persisted, or been repurposed because of their capacity to engage bacterial-like membranes or bacterial physiology. If naturally generated through proteolysis or other processing, such encrypted fragments could represent a previously unrecognized route by which mitochondrial genomes contribute to host immunity.

Testing this idea requires a way to convert evolutionary sequence information into experimentally tractable molecules. The emerging field of molecular de-extinction has shown that machine learning can identify antimicrobial candidates from extinct genomes and proteomes, with tools such as panCleave^8^ and APEX^9,10^ coupling proteolytic release of encrypted peptides with activity prediction. However, most de-extinction efforts have focused on organismal proteomes, leaving compact organellar genomes largely unexplored.

Here we test the biochemical premise that ancient human mtDNA harbors encrypted peptide sequences with intrinsic antimicrobial function. We combine archaeogenomic sequence data, ORF discovery, *in silico* proteolysis, and deep-learning prediction to identify mitochondria-derived encrypted peptides, which we call mitochondrins. By synthesizing and experimentally testing prioritized candidates, we show that mitochondrial coding space yields antibacterial peptides with motif-dependent activity, mechanistic diversity, selective activity over mammalian cells for some sequences, and initial efficacy in vivo. Several active peptides are also conserved in modern human mtDNA. Although our experiments do not establish endogenous production or a physiological immune role, they provide a direct biochemical foundation for the hypothesis that mitochondrial genomes may contribute to host defense.

## Results

### Ancient mitochondrial genomes reveal a constrained antimicrobial peptide landscape

To test whether ancient mitochondrial genomes encode latent antimicrobial peptides, we built a computational discovery pipeline that combined ORF extraction, in silico proteolysis, and APEX-based activity prediction (**Fig. 1a**). Screening 2,025 ancient human mtDNA sequences from AmtDB yielded 741,575 unique peptide substrings, candidate mitochondrins. APEX prioritized 65 candidates with predicted median MIC values below 80 µmol l^-1^ across bacterial strains. The relatively low active-candidate yield compared with previous genome- or proteome-scale screens likely reflects the compactness and high sequence similarity of mitochondrial genomes ^11–16^, underscoring that this search space is small but information-rich. We synthesized 38 candidates for experimental validation, allowing us to move directly from ancient DNA mining to functional testing.

**Fig. 1.**
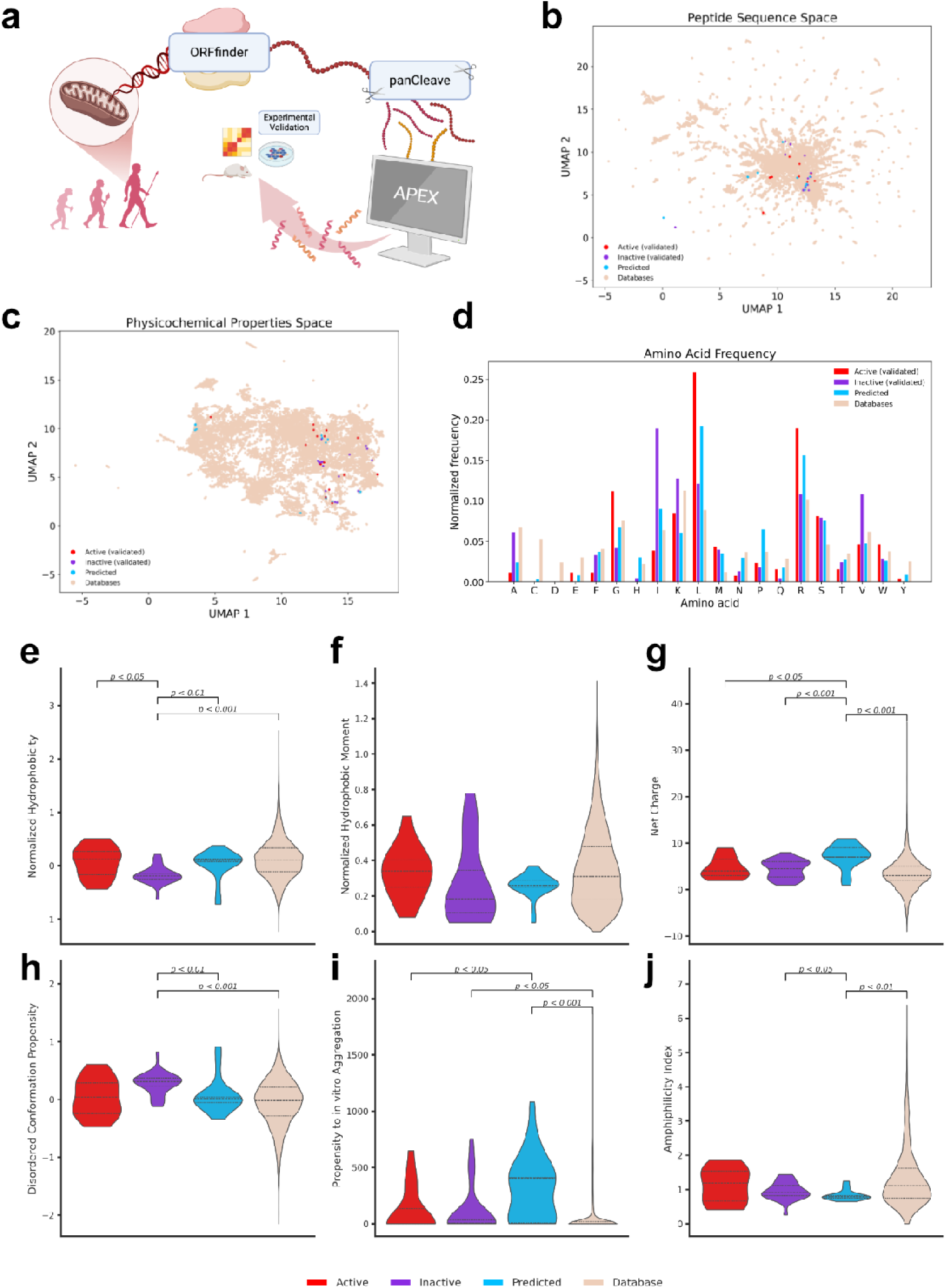
Ancient mitochondrial genomes encode a constrained peptide landscape with AMP-like features. **(a)** Schematic of the discovery workflow. ORFfinder identified open reading frames from ancient human mtDNA, panCleave generated putative proteolytic fragments, and APEX prioritized candidates for antimicrobial activity. **(b)** UMAP projection of pairwise sequence-similarity space comparing predicted mitochondrins with database AMPs from DBAASP, APD3 and DRAMP 3.0. Predicted peptides, experimentally active peptides and inactive peptides are shown relative to database AMPs. **(c)** UMAP projection of physicochemical property space using the same color scheme. **(d)** Normalized amino acid frequencies across peptide groups. **(e-j)** Distributions of normalized hydrophobicity, normalized hydrophobic moment, net charge, disordered-conformation propensity, propensity to aggregate in vitro and amphiphilicity index. Statistical significance was assessed using Mann-Whitney tests with false-discovery-rate correction; p-values are shown above plots.

To determine whether mitochondrins occupy known or underexplored regions of AMP space, we compared the 65 predicted peptides with 20,762 database AMPs (DBAASP^17^, APD3^18^, and DRAMP 3.0^19^). Pairwise sequence similarity followed by UMAP revealed that mitochondrins broadly overlap with known AMPs while also extending toward the periphery of the database-defined landscape (**Fig. 1b**). Active and inactive mitochondrins did not separate into obvious sequence clusters, indicating that activity is not captured by global sequence similarity alone. Instead, the distribution suggests that mitochondrial peptides combine recognizable AMP-like features with sequence constraints imposed by their organellar origin.

We next mapped physicochemical property space using DBAASP descriptors and UMAP (**Fig. 1b**). Mitochondrins again occupied regions overlapping with known AMPs but were enriched at the edges of the curated AMP landscape. This pattern is important because it indicates that ancient mtDNA does not simply recapitulate known AMP databases. Rather, it provides a compact, evolutionarily constrained source of peptides that resemble AMPs enough to be discoverable by machine learning while retaining properties that are underrepresented in existing collections.

At the amino-acid level, mitochondrins were enriched in leucine and arginine relative to database peptides (**Fig. 1d**). This compositional bias is consistent with mitochondrial biology, where leucine-rich hydrophobic segments and arginine-rich motifs contribute to protein stability, targeting, import and signaling. In contrast, database AMPs showed higher frequencies of several small or polar residues, including alanine, serine, aspartate and glutamate. Mitochondrins also contained fewer aromatic residues than curated AMPs, suggesting that some candidates may rely less on aromatic membrane anchoring and more on hydrophobic-cationic organization or electrostatic engagement. Together, these features define a mitochondrial AMP-like space that is constrained, recognizable and distinct.

Physicochemical comparisons reinforced this conclusion (**Fig. 1e-j**). Relative to database AMPs, mitochondrins showed narrower distributions of normalized hydrophobicity, hydrophobic moment, amphiphilicity and net charge, consistent with origin from a compact organellar genome rather than diverse natural AMP families. They also displayed higher predicted conformational order and greater variability in aggregation propensity. Although the experimentally tested set is limited, the repeated deviations from curated AMP distributions support a central point: mitochondrins are not random fragments of mtDNA, nor simple copies of known AMPs. They represent a distinct peptide landscape shaped by mitochondrial sequence constraints and accessible through computational mining.

### Evolutionary filtering prioritizes conserved mitochondrins

Because ancient DNA datasets can contain degradation, amplification, and reconstruction artifacts, we assessed the evolutionary support for experimentally validated mitochondrins using both the vertebrate mitochondrial genetic code and the standard genetic code. Among the active peptides, four - mitochondrins 11, 32, 39, and 40 - showed perfect sequence conservation when compared with modern human mtDNA^20^ under the vertebrate mitochondrial code. Mitochondrin- 32 was especially notable: it was conserved across ancient and modern comparisons and was the most broadly active peptide in the panel. This combination of conservation and potency suggests that at least some mitochondrins derive from robust mitochondrial coding information rather than from ancient-DNA noise. By contrast, eight peptides matched modern sequences only under the standard genetic code, consistent with codon-assignment effects during computational translation, and four peptides lacked modern matches under either code, which may reflect ancient-DNA artifacts or genuinely rare historical variation. These results provide an evolutionary filter for distinguishing high-confidence mitochondrial peptides from lower- confidence candidates ^21^. The conservation of active mitochondrins in modern human mtDNA is especially relevant to the host-defense hypothesis, because it shows that the encoded antibacterial potential is not restricted to ancient sequence variation.

### Mitochondrins display antimicrobial activity and motif-level sequence rules

We next asked whether computationally prioritized mitochondrins inhibit bacterial growth. We synthesized a diverse subset of 38 candidates and measured MICs against a panel of clinically relevant Gram-negative and Gram-positive pathogens (**Fig. 2a**), including drug-resistant strains (**Fig. 2a**). Seventeen peptides showed measurable antimicrobial activity, demonstrating that ancient mtDNA mining can yield experimentally validated antibacterial molecules. The activity profiles also provided an opportunity to infer sequence-activity rules directly from related mitochondrin families.

**Fig. 2.**
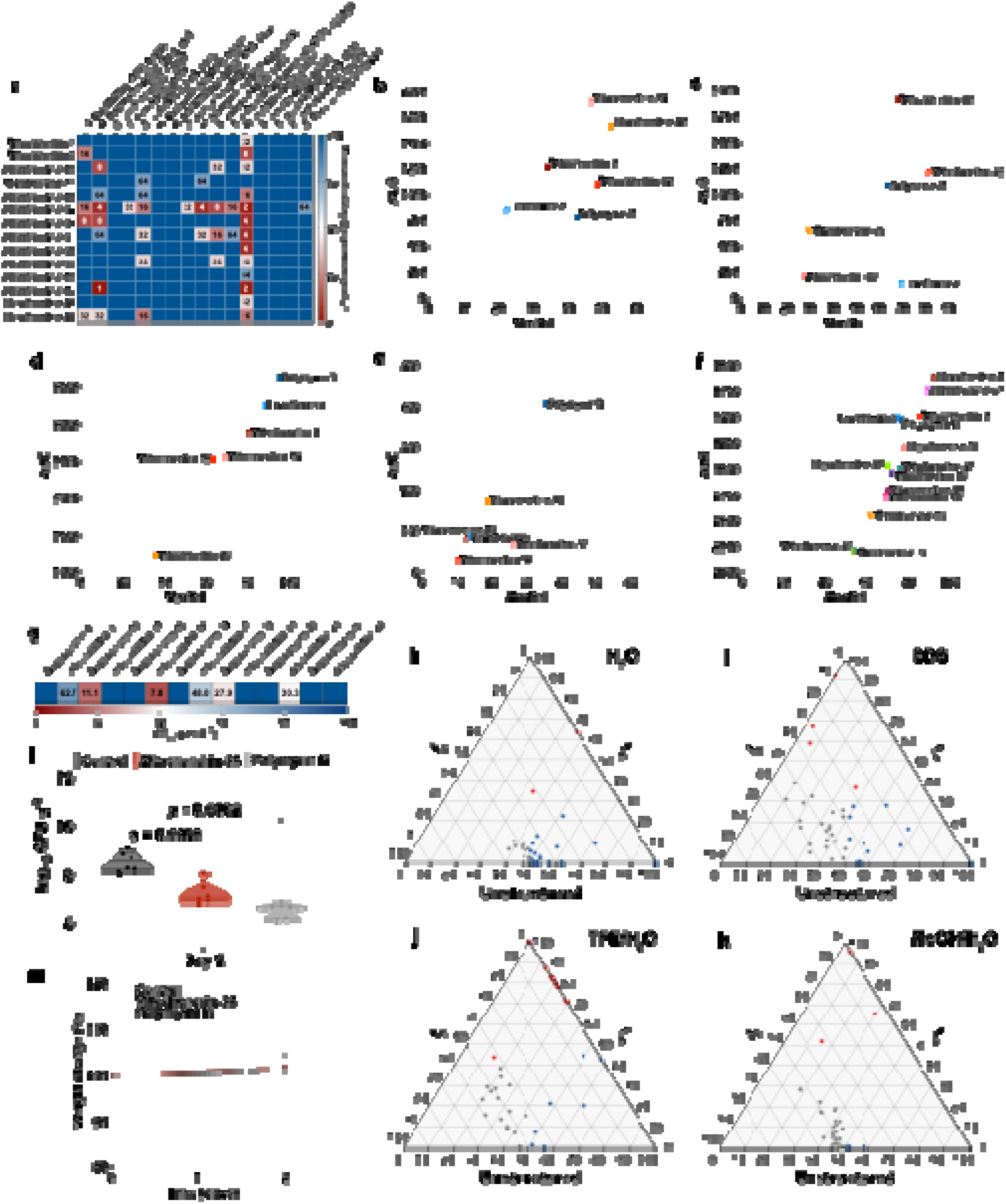
Antimicrobial activity, mechanism of action, and cytotoxic profiles of mitochondrins. **(a)** Heat map displaying the antimicrobial activities (μmol L^-1^) of mitochondrins from ancient mitochondria against clinically relevant pathogens, including antibiotic-resistant strains. Briefly, bacterial cells were incubated with serially diluted peptides (0-64 μmol L^-1^) at 37 °C. Bacterial growth was assessed by measuring the optical density at 600 nm in a microplate reader one day post-treatment. The MIC values presented in the heat map represent the mode of the replicates for each condition. To assess whether mitochondrins act on bacterial membranes, all active peptides against **(b)** *A. baumannii* ATCC 19606 and **(c)** *S. enterica* ATCC 9150 were subjected to outer membrane permeabilization and **(d)** *A. baumannii* ATCC 19606, **(e)** *S. enterica* ATCC 9150, and **(f)** *B. subtilis* ATCC 23857 were used in cytoplasmic membrane depolarization assays. The fluorescent probe 1-(N-phenylamino)naphthalene (NPN) was used to assess membrane permeabilization induced by the tested mitochondrins. The fluorescent probe 3,3′-dipropylthiadicarbocyanine iodide (DiSC_3_-5) was used to evaluate membrane depolarization caused by mitochondrins. The values displayed represent the fluorescence of both probes relative to untreated controls. **(g)** Cytotoxic concentrations against human embryonic kidney (HEK293T) cells leading to 50% cell lysis (CC_50_) were determined by interpolating the dose-response data using a non-linear regression curve. All experiments were performed in three independent replicates. Ternary plots showing the percentage of secondary structure for each peptide (at 50 μmol L^-1^) in four different solvents: **(h)** water, **(i)** sodium dodecyl sulfate (SDS, 10 mmol L^-1^) in water, **(j)** 60% trifluoroethanol (TFE) in water, and **(k)** 50% methanol in water. Secondary structure fractions were calculated using the BeStSel server^28^. Red dots indicate retrovirusins with higher helical content, while blue dots represent retrovirusins with higher β-content, and in gray peptides that were unstructured. **(l)** Anti-infective activity against *A. baumannii* ATCC 19606 in a skin abscess model was evaluated after topical peptide administration (n = 6). Two days after injection, mitochondrin-34 at 10×MIC exhibited a bactericidal effect when compared to the untreated control group. Polymyxin B was used as positive control. The limit of detection (LOD) for the CFU quantification is log_10_ CFU = 2. **(m)** Mouse weight was monitored throughout the duration of the skin abscess model to assess potential toxic effects of both the bacterial load and the mitochondrin. Statistical significance in panel **l** was determined using one- way ANOVA followed by Dunnett’s test; p-values are shown in the graphs. In the violin, the center line represents the mean, the box limits the first and third quartiles, and the whiskers (minima and maxima) represent 1.5 × the interquartile range. The solid line inside each box represents the mean value obtained for each group.

The MIC data revealed that mitochondrin activity is governed by motif architecture rather than by simple peptide length, charge, or hydrophobicity. Several candidates form nested families in which one peptide is a truncation or extension of another, creating internal controls for testing how small sequence changes influence potency.

One example is the MLSKLVNWKLTV(L) family (mitochondrins 2, 4, and 40). These peptides differ by only one to three terminal residues and share an almost identical hydrophobic and amphipathic core. Yet only mitochondrin-40 (MLSKLVNWKLTV) showed detectable activity, and only against *B. subtilis* at 64 µmol l^-1^, whereas the longer analog mitochondrin-2 (MLSKLVNWKLTVL) and the shorter analogs were inactive at the highest concentration tested. Thus, truncation alone does not unlock activity; instead, even modest terminal changes can determine whether a mitochondrial fragment crosses the threshold into antibacterial function.

A second nested series built around the VIIISSKARRVLIIKIKAKF motif (mitochondrins 20, 21, 22, 23, 24, 25, 26, and 35) further illustrates this principle. Despite a dense Lys/Arg-rich core and a hydrophobic segment, none of these peptides inhibited the tested strains at ≤64 µmol l^-1^. Progressive extension or shortening of this highly cationic scaffold therefore failed to generate activity, showing that positive charge and hydrophobicity are insufficient without the correct spatial patterning.

By contrast, the most active peptides, including mitochondrin-32 (LIGGWGLGWSGKRLRKILRRKKLLR) and mitochondrin-43 (SLSLMLTLIRGLSKRLG) appear as complete motifs rather than simple truncations. Both combine a hydrophobic or aromatic N-terminal region with a compact C-terminal cationic cluster. Mitochondrin-32 inhibited ten strains, with MICs as low as 2 µmol l^-1^ against *B. subtilis* and 4 µmol l^-1^ against *A. baumannii* ATCC BAA-1605 and *P. aeruginosa* ATCC BAA-3197. Mitochondrin-43 reached 1– 2 µmol l^-1^ against *A. baumannii* ATCC BAA-1605 and *B. subtilis*. Within the related SLSL family, mitochondrins 34 and 43 have the same length but differ by a few central substitutions, yet their potencies range from weak Gram-positive activity to broad low-micromolar activity. These comparisons show that mitochondrins encode high-resolution sequence rules in which motif placement, not overall composition, determines function.

Taken together, the antimicrobial data transform mitochondrins from predicted fragments into experimentally validated sequence families. The nested scaffolds serve as natural structure-activity probes: most related variants remain inactive, whereas specific hydrophobic-cationic configurations produce potent activity. This motif-level resolution is a key advantage of mining compact organellar genomes, where related fragments can reveal the sequence logic of activity. These data therefore establish sequence-encoded antibacterial potential within mitochondrial coding space, a prerequisite for any possible contribution to host defense.

### Mitochondrins engage bacterial membranes through different mechanisms

To understand how active mitochondrins affect bacteria, we measured outer-membrane permeabilization with NPN and cytoplasmic membrane depolarization with DiSC_3_-5 in *A. baumannii* ATCC 19606, *B. subtilis* ATCC 23857, and *S. enterica* ATCC 9150 (**Fig. 2b-f**). These assays revealed that mitochondrins do not behave as a single class of uniformly lytic peptides. Instead, they engage bacterial envelopes through species-dependent and sequence-dependent mechanisms.

In NPN uptake assays, some mitochondrins produced large and sustained fluorescence increases, consistent with strong outer-membrane permeabilization, whereas others generated weak, transient or biphasic responses. This divergence was observed in both *A. baumannii and S. enterica* (**Fig. 2b,c**). Notably, several peptides with strong MIC activity caused little detectable NPN uptake, indicating that potent growth inhibition does not necessarily require extensive outer-membrane disruption.

DiSC_3_-5 assays further separated the mitochondrins by their ability to dissipate cytoplasmic membrane potential. In *A. baumannii*, some peptides induced sustained depolarization, whereas others produced only modest signals (**Fig. 2d**). In *S. enterica*, depolarization was more selective, with only a subset of active peptides producing clear responses above baseline (**Fig. 2e**). In *B. subtilis*, responses spanned a continuum from rapid depolarization to minimal perturbation (**Fig. 2f**).

Integrating both assays revealed at least three functional classes: peptides that strongly permeabilize and depolarize membranes; peptides that induce moderate or transient envelope stress; and peptides that retain antimicrobial potency despite weak membrane readouts. The absence of a strict correlation between MIC potency and membrane disruption suggests that some mitochondrins act through classical membrane-active mechanisms, whereas others may combine subtle envelope perturbation with intracellular or multi-target effects.

This mechanistic diversity tracks with sequence architecture. Strongly lytic mitochondrins generally combine extended hydrophobic or aromatic segments with balanced cationic domains, consistent with amphipathic membrane insertion. Peptides that kill with weaker membrane signatures often contain compact Arg/Lys clusters and differ from inactive relatives by only a few residues. Thus, ancient mtDNA appears to encode not only antimicrobial potency, but a tunable repertoire of bacterial-envelope interactions.

### Mitochondrins can combine potency with low mammalian-cell toxicity

We next evaluated whether mitochondrin activity against bacteria is separable from toxicity toward mammalian cells. HEK293T cytotoxicity assays showed a broad range of safety profiles (**Fig. 2g**). Several peptides maintained high cell viability across the tested concentration range, whereas others reduced viability at lower concentrations, indicating narrower therapeutic windows.

Comparison of cytotoxicity and MIC values identified mitochondrins with favorable preliminary therapeutic indices. Particularly important are peptides that inhibit multiple Gram-negative strains while producing limited effects on HEK293T cells, because this profile suggests selective engagement of bacterial envelopes over mammalian membranes. Conversely, peptides with weak antimicrobial activity but greater cytotoxicity represent lower-priority starting points for therapeutic optimization.

Sequence features helped explain these differences. Less cytotoxic mitochondrins tended to distribute hydrophobic and cationic residues in balanced amphipathic patterns, whereas more cytotoxic peptides often contained dense hydrophobic segments or highly concentrated cationic motifs. This relationship paralleled the mechanistic assays: peptides with strong depolarization or pronounced NPN uptake often showed increased mammalian-cell effects, while peptides that killed bacteria with weaker membrane perturbation generally had milder cytotoxicity.

These observations are significant because they show that the mitochondrial peptide landscape contains both activity and selectivity. Mitochondrins therefore provide not only hits for antibacterial discovery but also sequence templates for optimization, where potency can be retained while membrane-aggressive features are tuned to reduce host-cell toxicity. The separation between antibacterial activity and mammalian-cell toxicity is particularly relevant to a host-defense model, in which endogenous effectors would need to preferentially engage microbial over host membranes.

### Active mitochondrins adopt multiple structural solutions

To determine whether ancient mitochondrins share a common structural basis for activity, we used circular dichroism and BeStSel deconvolution to analyze the 38 synthesized peptides in water and in membrane-mimicking environments (SDS/H_2_O, TFE/H_2_O and MeOH/H_2_O; **Figs. 2h-k and S1**). Most peptides were not strongly α-helical in aqueous solution. Instead, they showed mixed conformations with β-like and disordered components, indicating that many mitochondrins are only partially pre-organized before encountering membrane-like conditions.

Within the active subset, one structural class was clearly helix-inducible. Mitochondrins 34 and 43 transitioned from modest helicity in water to strong α-helical signatures in TFE/H_2_O, and mitochondrin-43 remained highly helical in SDS/H_2_O and MeOH/H_2_O. These data support a model in which precisely arranged hydrophobic and cationic motifs stabilize amphipathic helices upon membrane contact.

A second active class retained mixed or non-helical conformations even in membrane-mimicking media. Mitochondrin-32, the broadest-spectrum peptide, did not adopt a dominant α-helical state; instead, it preserved substantial β-antiparallel and unordered content across solvents. This structural profile is consistent with its mechanism-of-action data, in which potent antimicrobial activity occurred despite limited outer-membrane permeabilization and depolarization. Thus, some mitochondrins appear to act outside the canonical helix-driven membrane-disruption paradigm.

Inactive peptides reinforced the same conclusion. Several members of the VIIISSKARRVLIIKIKAKF-derived family became α-helical in TFE/H_2_O but remained inactive, and most MLSKLVNWKLTV(L) family members shared structural tendencies without measurable activity. Therefore, secondary structure alone does not classify mitochondrins as active or inactive. Antimicrobial function emerges from the intersection of sequence context, residue patterning, and inducible structure.

Cytotoxicity also did not map onto a single structural state. Mitochondrin-34 combined antimicrobial activity, helix induction and one of the most favorable cytotoxicity profiles in the panel. Mitochondrin-32 was highly potent but more cytotoxic despite lacking a dominant helical signature, whereas mitochondrin-43 showed strong helix stabilization with intermediate cytotoxicity. These comparisons indicate that mammalian-cell effects depend on how structure interacts with hydrophobic clustering, cationic density, and membrane selectivity.

Overall, the structural data reveal that mitochondrins converge on function without converging on one fold. Active peptides exploit at least two solutions: inducible amphipathic helices and mixed β-rich or disordered states compatible with antibacterial activity. This structural heterogeneity strengthens the central conclusion that ancient mitochondrial genomes encode a diverse, noncanonical repertoire of antimicrobial peptides.

### A mitochondrin reduces bacterial burden *in vivo*

To test whether mitochondrin activity extends beyond *in vitro* assays, we evaluated a representative active peptide in a murine skin abscess model of *A. baumannii* ATCC 19606 infection. This model provides a stringent first assessment of activity in tissue, where peptide stability, diffusion and host factors can limit efficacy.

At day 2 post-infection, a single topical dose of mitochondrin-34 at 10×MIC reduced bacterial burden relative to untreated controls (**Fig. 2l**). Polymyxin B produced a stronger reduction, as expected for an optimized reference antibiotic, but the mitochondrin effect demonstrates that ancient mtDNA-derived peptides can retain antibacterial activity in a complex biological setting. Mouse body weight remained stable throughout the experiment (**Fig. 2m**), supporting tolerability under the conditions tested. These results provide an initial *in vivo* proof of concept and motivate future optimization for stability, formulation and dosing.

## Discussion

This study supports a broader model of mitochondrial immunity in which the mitochondrial genome itself may harbor direct antimicrobial potential. By mining ancient human mtDNA, we identified encrypted peptides with measurable activity against Gram-negative and Gram-positive bacteria, diverse mechanisms of action, and, for some sequences, preliminary selectivity and activity in tissue. Although we do not yet know whether mitochondrins are naturally produced during infection, these findings make testable the hypothesis that mtDNA may contribute directly to host defense.

The rationale for this hypothesis is rooted in endosymbiosis. Mitochondria derive from a bacterial ancestor, retain bacterial-like molecular features, and are integrated into innate immune signaling. Mitochondrins could therefore arise through several non-mutually exclusive routes: retention of ancestral bacterial sequence motifs, host-driven adaptation related to control of the endosymbiont or competing microbes, or antimicrobial activity that emerges when mitochondrial proteins are fragmented. The perfect conservation of several active mitochondrins between ancient and modern human mtDNA is especially notable because it places this encoded antimicrobial potential within contemporary mitochondrial sequence space, rather than restricting it to ancient-DNA variation.

The experimental data argue that this activity is not explained by generic cationic peptide chemistry alone. Mitochondrins occupy a constrained physicochemical landscape, and closely related sequence families show that small changes in motif placement can switch antibacterial activity on or off. This high-resolution sequence dependence is important for the immune hypothesis because it suggests that antimicrobial function arises from specific encoded architectures rather than from arbitrary positively charged fragments.

Mechanistically, mitochondrins also extend beyond a single canonical antimicrobial-peptide model. Some strongly permeabilize and depolarize bacterial membranes, whereas others remain potent despite weak membrane signatures, pointing to more subtle envelope effects, intracellular targets, or multi-step mechanisms. Such diversity could be advantageous for host defense, where effectors encounter distinct bacterial species and physiological states.

Selectivity provides another important criterion for a potential immune effector. Several mitochondrins inhibit bacteria while producing comparatively limited effects on human cells, indicating that the mitochondrial peptide landscape contains sequences capable of distinguishing microbial from mammalian membranes.

The central limitation is whether mitochondrins are naturally produced, processed, and deployed by human cells. The present computational workflow predicts ORFs and active fragments but does not establish endogenous peptide abundance, subcellular localization, or infection-responsive release. Future studies should use targeted peptidomics and proteomics to detect mitochondrins in cells and tissues, determine whether their abundance changes during infection, inflammation, or mitochondrial stress, identify the proteases and processing pathways that generate them, and test whether genetic or pharmacological perturbation of candidate sequences alters antimicrobial phenotypes.

Taken together, by showing that sequences encoded in ancient human mitochondrial genomes possess antibacterial activity, we establish a biochemical foundation for testing whether mitochondria contribute directly to host defense through peptide effectors.

## Reporting summary

Further information on research design is available in the Nature Portfolio Reporting Summary linked to this article.

## Data and code availability

All original code for APEX 1.1 including ensemble weights, model architecture, AAindex1 feature table, and feature-construction utilities, has been deposited at GitLab (https://gitlab.com/machine-biology-group-public/apex-pathogen). PanCleave’s orginal code is also available at GitLab (https://gitlab.com/machine-biology-group-public/pancleave). Both are publicly available as of the date of publication. Any additional information required to reanalyze the data reported in this paper is available from the lead contact upon reasonable request.

## Acknowledgements

Cesar de la Fuente-Nunez holds a Presidential Professorship at the University of Pennsylvania. Research in this publication was supported by the National Institute of General Medical Sciences of the National Institutes of Health under award number R35GM138201 and the Defense Threat Reduction Agency (DTRA; HDTRA1-21-1-0014). We thank de la Fuente Lab members for insightful discussions. We also thank Benjamin Galeota-Sprung for providing insight into the evolutionary analysis of our mitochondrial peptides. Figures created with BioRender.com are attributed as such.

## Author contributions

M.D.T.T., A.A., F.W., and C.F.-N. conceptualized and designed the study. M.D.T.T. performed experiments and interpreted the data. H.-S.L. performed peptide synthesis. A.A. and F.W. performed the computational investigation and interpreted the data. All authors wrote and revised the manuscript.

## Methods

### Ancient mitochondrial DNA source

Ancient human mitochondrial DNA (mtDNA) sequences were obtained from the AmtDB website (https://amtdb.org/, access time: July 2024). These sequences represent a collection of ancient mtDNA samples obtained through archaeological studies. In total, 2,025 DNA sequences were downloaded as a single FASTA file for downstream computational processing.

### Translation of nucleotide sequences

Open reading frames (ORFs) were identified using the stand-alone version of NCBI’s ORFfinder (https://ftp.ncbi.nlm.nih.gov/genomes/TOOLS/ORFfinder/linux-i64/; Linux 64-bit). The minimum ORF threshold was set to 24 nucleotides. This value reflects a lower bound for functional AMP length, as β-sheets can span a membrane at ∼8 amino acids whereas α-helical peptides span a wider sequence length range which is usually >12 amino acid residues long (17).

### *In silico* proteolysis

The full-length translated peptides (741,575) were subjected to *in silico* proteolysis using panCleave, an in-house Python pipeline for simulating protease cleavage (https://gitlab.com/machine-biology-group-public/pancleave, access time: July 2024)^4^. This model predicts biologically relevant cleavage sites based on experimentally derived human protease substrates^4^. After running panCleave, 6,145 fragments were obtained. The peptides selected for chemical synthesis and experimental validation were all identified by APEX.

### Computational antimicrobial prediction

Peptide fragments were screened for antimicrobial activity using APEX1.1, a deep learning pipeline developed by our lab for classification of AMPs using regression modeling. The model was trained on curated sets of both non-AMPs and AMPs from both public and in-house datasets. Performance metrics and further description of model tuning are reported in Torres et al.^6^. Setting a maximum MIC median threshold of 80 μmol l^-1^, we obtained 65 unique peptide sequences. For experimental validation, 38 peptides were chosen considering potential for chemical synthesis and aggregation propensity.

### Physicochemical property analysis

Physicochemical properties were calculated using DBAASP’s tool^20^ using the Eisenberg and Weiss hydrophobicity scale^25^ for all peptides. The twelve properties included normalized hydrophobic moment, normalized hydrophobicity, net charge, isoelectric point, penetration depth, tilt angle, disordered conformation propensity, linear moment, propensity to *in vitro* aggregation, angle subtended by the hydrophobic residues, amphiphilicity index, and propensity to PPII coil.

### Peptide sequence-space visualization

For our peptide dataset, we performed pairwise sequence-similarity comparison to generate a similarity matrix. We then applied UMAP to reduce the sequence-similarity space to two dimensions and visualize the resulting peptide sequence landscape.

### Evolutionary validation

The BLAST+ executable (https://ftp.ncbi.nlm.nih.gov/blast/executables/blast+/LATEST/, version 2.17.0+)^26^ was used to compare 17 validated peptides to ancient (AmtDB) and modern (MitoMap) mitochondrial DNA sources. The tblastn command was employed with specified parameters to better account for our short sequences: word size 2, PAM30 matrix, E-value of 1000. Searches were conducted with both the vertebrate mitochondrial table (-db_gencode 2), and the standard genetic code (-db_gencode 1) to highlight the significance of proper code selection. Sequences with a 100% sequence identity match against the mitochondrial code were classified as evolutionarily conserved.

### Peptide synthesis

Peptides were synthesized on an automated peptide synthesizer (Symphony X, Gyros Protein Technologies) by standard Fmoc-based solid-phase peptide synthesis (SPPS) on Fmoc-protected amino acid–Wang resins (100–200 mesh). N,N-Dimethylformamide (DMF) was used as the primary solvent throughout synthesis. Stock solutions included: 500 mmol L^-1^ Fmoc-protected amino acids in DMF, a coupling mixture of HBTU (450 mmol L^-1^) and N-methylmorpholine (NMM, 900 mmol L^-1^) in DMF, and 20% (v/v) piperidine in DMF for Fmoc deprotection. After synthesis, peptides were deprotected and cleaved from the resin using a cleavage cocktail of trifluoroacetic (TFA)/triisopropylsilane (TIS)/dithiothreitol (DTT)/water (92.8% v/v, 1.1% v/v, 0.9% w/v, 4.8%, w/w) for 2.5 hours with stirring at room temperature. The resin was removed by vacuum filtration, and the peptide-containing solution was collected. Crude peptides were precipitated with cold diethyl ether and incubated for 20 min at -20 °C, pelleted by centrifugation, and washed once more with cold diethyl ether. The resulting pellets were dissolved in 0.1% (v/v) aqueous formic acid and incubated overnight at -20 °C, followed by lyophilization to obtain dried peptides. For characterization, peptides were dried, reconstituted in 0.1% formic acid, and quantified spectrophotometrically. Peptide separations were performed on a Waters XBridge C_18_ column (4.6 × 50 mm, 3.5 µm, 120 Å) at room temperature using a conventional high-performance liquid chromatography (HPLC) system. Mobile phases were water with 0.1% formic acid (solvent A) and acetonitrile with 0.1% formic acid (solvent B). A linear gradient of 1–95% B over 7 min was applied at 1.5 mL min^-1^. UV detection was monitored at 220 nm. Eluates were analyzed on Waters SQ Detector 2 with electrospray ionization in positive mode. Full scan spectra were collected over m/z 100–2,000. Selected Ion Recording (SIR) was used for targeted peptides. Source conditions were capillary voltage 3.0 kV, cone voltage 25-40 V, source temperature 120 °C, and desolvation temperature 350 °C. Mass spectra were processed with MassLynx software. Observed peptide masses were compared with theoretical values, and quantitative analysis was based on integrated SIR peak areas.

### Bacterial strains and growth conditions

The bacterial panel included the following strains: *Acinetobacter baumannii* ATCC 19606; *A. baumannii* ATCC BAA-1605 (resistant to ceftazidime, gentamicin, ticarcillin, piperacillin, aztreonam, cefepime, ciprofloxacin, imipenem, and meropenem); *Escherichia coli* ATCC 11775; *E. coli* ATCC BAA-3170 (resistant to colistin and polymyxin B); *Enterobacter cloacae* ATCC 13047; *Klebsiella pneumoniae* ATCC 13883; *K. pneumoniae* ATCC BAA-2342 (resistant to ertapenem and imipenem); *Pseudomonas aeruginosa* PAO1; *P. aeruginosa* ATCC BAA-3197 (resistant to fluoroquinolones, β-lactams, and carbapenems); *Salmonella enterica* ATCC 9150; *S. enterica* subsp. *enterica* Typhimurium ATCC 700720; *Bacillus subtilis* ATCC 23857; *Staphylococcus aureus* ATCC 12600; *S. aureus* ATCC BAA-1556 (methicillin-resistant); *Enterococcus faecalis* ATCC 700802 (vancomycin-resistant); and *Enterococcus faecium* ATCC 700221 (vancomycin-resistant). *P. aeruginosa* strains were propagated on Pseudomonas Isolation Agar, whereas all other species were maintained on Luria-Bertani (LB) agar and broth. For each assay, cultures were initiated from single colonies, incubated overnight at 37 °C, and subsequently diluted 1:100 into fresh medium to obtain cells in mid-logarithmic phase.

### Minimal inhibitory concentration determination

Broth microdilution assays^27^ were performed to determine the minimum inhibitory concentration (MIC) values of each peptide. Peptides were added to nontreated polystyrene microtiter 96-well plates and 2-fold serially diluted in sterile water from 1 to 64 μmol L^-1^. Bacterial inoculum at 4×10^6^ CFU mL^-1^ in LB medium was mixed 1:1 with the peptide. The MIC was defined as the lowest concentration of peptide able to completely inhibit the bacterial growth after 24 h of incubation at 37 °C. All assays were done in three independent replicates.

### Outer-membrane permeabilization assays

NPN uptake assays were used to assess outer-membrane permeabilization. Inocula of *A. baumannii* ATCC 19606 and *S. enterica* ATCC 9150 were grown to an OD at 600 nm of 0.4 mL^-1^, centrifuged (9,391 ×g for 3 min), washed and resuspended in 5 mmol L^-1^ HEPES buffer (pH 7.4) containing 5 mmol L^-1^ glucose. The bacterial solution was added to a white 96-well plate (100 μL per well) together with 4 μL of NPN at 0.5 mmol L^-1^. Consequently, peptides diluted in water were added to each well, and the fluorescence was measured at λ_ex_ = 350 nm and λ_em_ = 420 nm over time for 45 min. The relative fluorescence was calculated using the untreated control (buffer + bacteria + fluorescent dye) as baseline and the following equation was applied to reflect % of difference between the baselines and the sample:

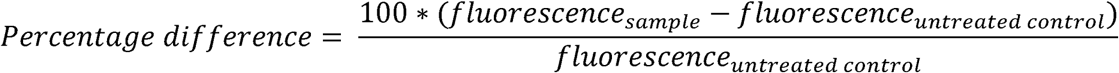

### Cytoplasmic membrane depolarization assays

Membrane depolarization was measured with DiSC_3_-5. Mid-log-phase *A. baumannii* ATCC 19606, *S. enterica* ATCC 9150, and *B. subtilis* ATCC 23857 were washed (9,391 ×g for 3 min) and resuspended at 0.05 OD mL^-1^ (optical value at 600 nm) in HEPES buffer (pH 7.2) containing 20 mmol L^-1^ glucose and 0.1 mol L^-1^ KCl. DiSC_3_-5 at 20 μmol L^-1^ was added to the bacterial suspension (100 μL per well) for 15 min to stabilize the fluorescence which indicates the incorporation of the dye into the bacterial membrane, and then the peptides were mixed 1:1 with the bacteria to a final concentration corresponding to their MIC_100_ values. Membrane depolarization was then followed by reading changes in the fluorescence (λ_ex_ = 622 nm, λ_em_ = 670 nm) over time for 60 min. The relative fluorescence was calculated using the untreated control (buffer + bacteria + fluorescent dye) as baseline and the following equation was applied to reflect % of difference between the baselines and the sample:

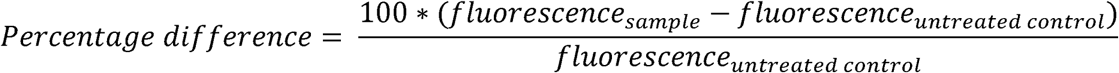

### Cytotoxicity assays

The cells were cultured in high-glucose Dulbecco’s modified Eagle’s medium supplemented with 1% penicillin and streptomycin (antibiotics) and 10% fetal bovine serum and grown at 37 °C in a humidified atmosphere containing 5% CO_2_.

One day prior to the experiment, 100 μL aliquots of human embryonic kidney (HEK293T) cells, at a concentration of 50,000 cells per mL, were seeded into each well of 96-well plates (5,000 cells per well). Following cell attachment, the HEK293T cells were treated with increasing concentrations of peptides (ranging from 8 to 128 μmol L^-1^) and incubated for 24 h. After the exposure period, cytotoxicity was assessed using the 3-(4,5-dimethylthiazol-2- yl)-2,5-diphenyltetrazolium bromide (MTT) assay. Specifically, the MTT reagent was prepared at a concentration of 0.5 mg mL^-1^ in phenol red-free medium and used to replace the peptide-containing supernatants (100 μL per well). The plates were then incubated for 4 h at 37 °C in a humidified atmosphere with 5% CO, facilitating the formation of insoluble formazan crystals. These crystals were subsequently dissolved in 0.04 mol L^-1^ hydrochloric acid prepared in anhydrous isopropanol. Absorbance was measured at 570 nm using a spectrophotometer to quantify cell viability. All experiments were conducted in triplicate (three biological replicates).

### Circular dichroism experiments

The circular dichroism experiments were conducted using a J1500 circular dichroism spectropolarimeter (Jasco) in the Biological Chemistry Resource Center (BCRC) at the University of Pennsylvania. Experiments were performed at 25 °C, the spectra graphed are an average of three accumulations obtained with a quartz cuvette with an optical path length of 1.0 mm, ranging from 260 to 190 nm at a rate of 50 nm min^-1^ and a bandwidth of 0.5 nm. The concentration of all peptides tested was 50 μmol L^-1^, and the measurements were performed in water, a mixture of trifluoroethanol (TFE) and water in a 3:2 ratio, a mixture of methanol (MeOH) and water in a 1:1 ratio, and sodium dodecyl sulfate (SDS) in water at 10 mmol L^-1^, with respective baselines recorded prior to measurement. A Fourier transform filter was applied to minimize background effects. Secondary structure fraction values were calculated using the single spectra analysis tool on the server BeStSel.

### Skin abscess infection mouse model

The backs of anesthetized six-week-old female CD-1 mice were shaved and injured with a superficial linear skin abrasion made with a needle. An aliquot of *A. baumannii* ATCC 19606 (9.0×10^5^ CFU mL^-1^; 20 μL) previously grown in LB medium until 0.5 OD mL^-1^ (optical value at 600 nm) and then washed twice with sterile PBS (pH 7.4, 9,391 ×g for 3 min) was added to the scratched area. Peptides diluted in sterile water at their MIC value were administered to the wounded area 1 h post-infection. Two days post-infection, animals were euthanized, and a uniform section of scarified skin was excised, homogenized using a bead beater (25 Hz for 20 min), 10-fold serially diluted, and plated on McConkey agar plates for CFU quantification. The experiments were performed using six mice per group. Mice were single-housed to avoid cross-contamination and maintained under a 12-hour light/dark cycle at 22 °C with humidity controlled at 50%. The skin abscess infection mouse model was revised and approved by the University Laboratory Animal Resources (ULAR) from the University of Pennsylvania (Protocol 806763).

---

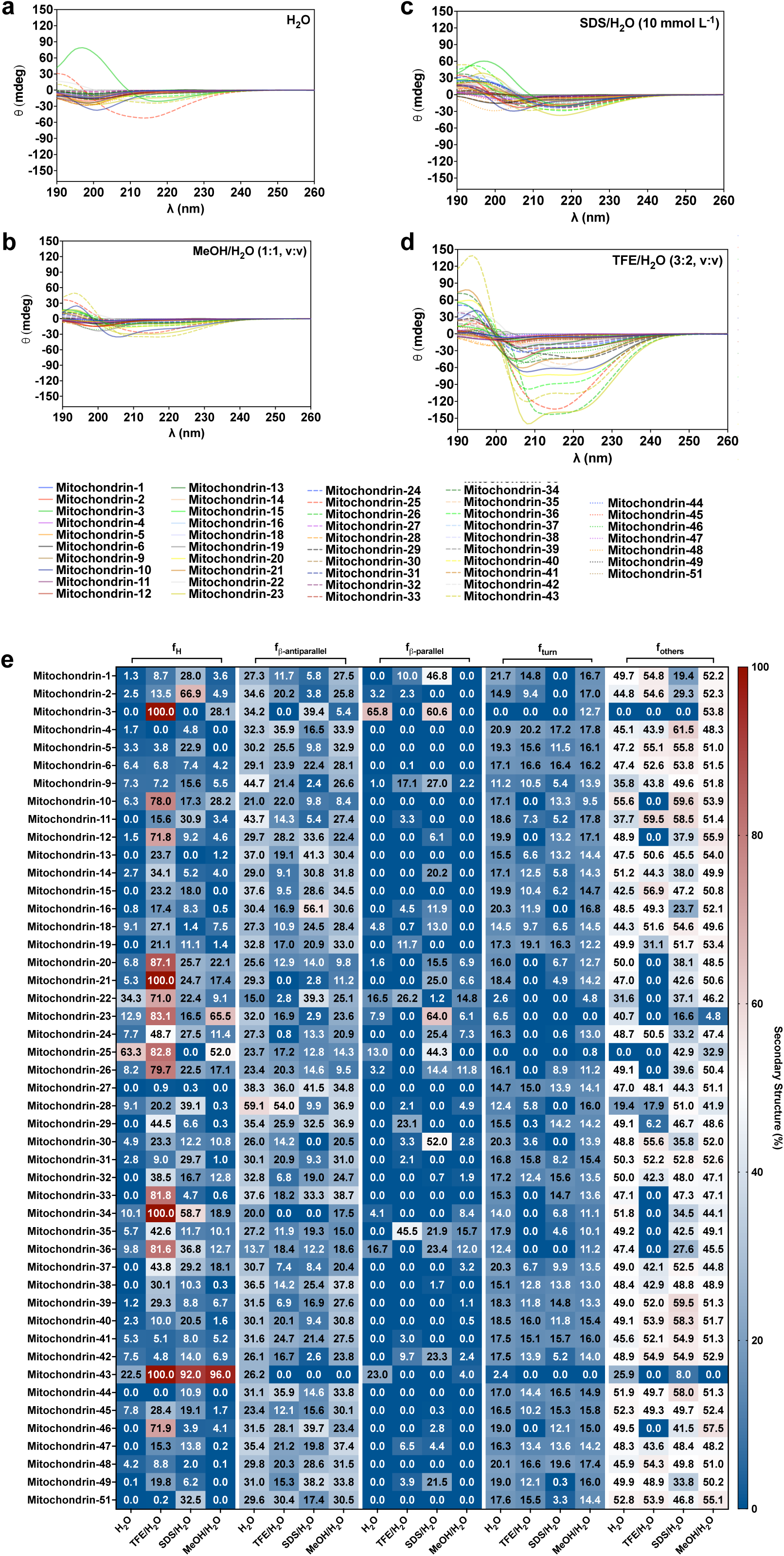

## References

1. Zimorski, V., Ku, C., Martin, W. F. & Gould, S. B. Endosymbiotic theory for organelle origins. Curr. Opin. Microbiol. 22, 38–48 (2014).

2. Gray, M. W. Mitochondrial Evolution. Cold Spring Harb. Perspect. Biol. 4, a011403– a011403 (2012).

3. Bennett, G. M., Kwak, Y. & Maynard, R. Endosymbioses Have Shaped the Evolution of Biological Diversity and Complexity Time and Time Again. Genome Biol. Evol. 16, (2024).

4. Marques, E., Kramer, R. & Ryan, D. G. Multifaceted mitochondria in innate immunity. npj Metabolic Health and Disease 2, 6 (2024).

5. Liu, X.-Y., Wei, B., Shi, H.-X., Shan, Y.-F. & Wang, C. Tom70 mediates activation of interferon regulatory factor 3 on mitochondria. Cell Res. 20, 994–1011 (2010).

6. Seth, R. B., Sun, L., Ea, C.-K. & Chen, Z. J. Identification and Characterization of MAVS, a Mitochondrial Antiviral Signaling Protein that Activates NF-κB and IRF3. Cell 122, 669–682 (2005).

7. Li, D. & Wu, M. Pattern recognition receptors in health and diseases. Signal Transduct. Target. Ther. 6, 291 (2021).

8. Maasch, J. R. M. A., Torres, M. D. T., Melo, M. C. R. & de la Fuente-Nunez, C. Molecular de-extinction of ancient antimicrobial peptides enabled by machine learning. Cell Host Microbe 31, 1260–1274 (2023).

9. Wan, F., Torres, M. D. T., Peng, J. & de la Fuente-Nunez, C. Deep-learning-enabled antibiotic discovery through molecular de-extinction. *Nat*. Biomed. Eng. 8, 854–871 (2024).

10. Torres, M. D. T., Wan, F. & de la Fuente-Nunez, C. Deep learning reveals antibiotics in the archaeal proteome. Nat. Microbiol. 2153–2167 (2025) doi:10.1038/s41564-025-02061-0.

11. Torres, M. D. T. et al. Mining for encrypted peptide antibiotics in the human proteome. Nat. Biomed. Eng. 6, 67–75 (2022).

12. Torres, M. D. T., Cesaro, A. & de la Fuente-Nunez, C. Peptides from non-immune proteins target infections through antimicrobial and immunomodulatory properties. Trends Biotechnol. (2024) doi:10.1016/j.tibtech.2024.09.008.

13. Xia, X., Torres, M. D. T. & de la Fuente-Nunez, C. Proteasome-derived antimicrobial peptides discovered via deep learning. bioRXiv (2025) doi:10.1101/2025.03.17.643752.

14. Guan, C., Torres, M. D. T., Li, S. & de la Fuente-Nunez, C. Computational exploration of global venoms for antimicrobial discovery with Venomics artificial intelligence. Nat. Commun. 16, 6446 (2025).

15. Torres, M. D. T. et al. Mining human microbiomes reveals an untapped source of peptide antibiotics. Cell 187, 5453–5467 (2024).

16. Santos-Júnior, C. D. et al. Discovery of antimicrobial peptides in the global microbiome with machine learning. Cell 187, 3761–3778 (2024).

17. Pirtskhalava, M. et al. DBAASP v3: database of antimicrobial/cytotoxic activity and structure of peptides as a resource for development of new therapeutics. Nucleic Acids Res. 49, D288–D297 (2021).

18. Wang, G., Li, X. & Wang, Z. APD3: the antimicrobial peptide database as a tool for research and education. Nucleic Acids Res. 44, D1087–D1093 (2016).

19. Shi, G. et al. DRAMP 3.0: an enhanced comprehensive data repository of antimicrobial peptides. Nucleic Acids Res. 50, D488–D496 (2022).

20. Osawa, S., Ohama, T., Jukes, T. H. & Watanabe, K. Evolution of the mitochondrial genetic code I. Origin of AGR serine and stop codons in metazoan mitochondria. J. Mol. Evol. 29, 202–207 (1989).

21. Parr, R. L. et al. The pseudo-mitochondrial genome influences mistakes in heteroplasmy interpretation. BMC Genomics 7, 185 (2006).

