## Supplementary Information for "Ancient human mitochondrial genomes encode antimicrobial peptides"

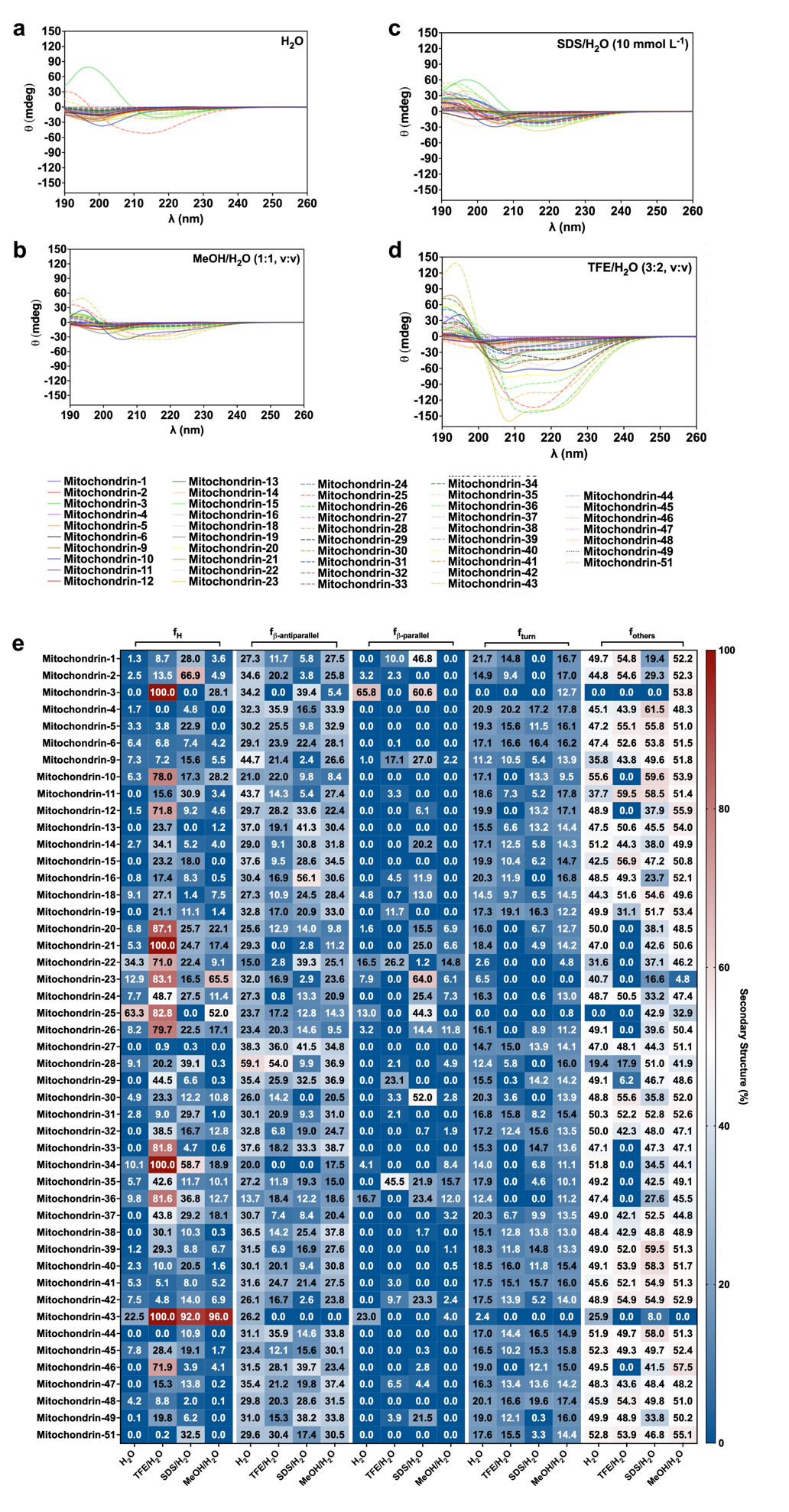

**Fig. S1.** **Circular dichroism spectra of mitochondrins.** Circular dichroism experiments were conducted with peptides from the retroviruses using a J-1500 Jasco circular dichroism spectrophotometer. The spectra were recorded in four different media: **(a)** water, **(b)** 50% methanol in water, **(c)** Sodium dodecyl sulfate (SDS) in water (10 mmol L^-1^), and **(d)** 60% trifluoroethanol in water, after three accumulations at 25 ^o^C, using a 1 mm path length quartz cell, between 260 and 190 nm at 50 nm min^-1^, with a bandwidth of 0.5 nm. The concentration of all peptides tested was 50 μmol L^-1^. **(d)** Heatmap with the percentage of secondary structure found for each peptide in the same solvents. Secondary structure fraction was calculated using the BeStSel server.
